# Single-Round Remodeling of the Active Site of an Artificial Metalloenzyme using an Ultrahigh-Throughput Double Emulsion Screening Assay

**DOI:** 10.1101/2021.09.20.460989

**Authors:** Jaicy Vallapurackal, Ariane Stucki, Alexandria Deliz Liang, Juliane Klehr, Petra S. Dittrich, Thomas R. Ward

**Affiliations:** Department of Chemistry, University of Basel, Mattenstrasse 24a, 4058 Basel, Switzerland; Department of Biosystems Science and Engineering, ETH Zürich, Mattenstrasse 24a, 4058 Basel, Switzerland; NCCR Molecular Systems Engineering, 4058 Basel, Switzerland

## Abstract

The potential of high-throughput compartmentalization renders droplet microfluidics an attractive tool for directed evolution of enzymes as it permits maintenance of the phenotype-genotype linkage throughout the entire optimization procedure. In particular, water-in-oil-in-water double emulsions droplets (DEs) produced by microfluidics enable the analysis of reaction compartments at ultra-high-throughput using commercially available fluorescence-activated cell sorting (FACS) devices. Here we report a streamlined method applicable for the ultrahigh-throughput screening of an artificial metalloenzyme (ArM), an artificial deallylase (ADAse), in double emulsions. The DE-protocol was validated by screening a four hundred member, double-mutant streptavidin library for the CpRu-catalyzed uncaging of aminocoumarin. The most active variants, identified by next generation sequencing of the sorted DE droplets with highest fluorescent intensity, are in good agreement with 96-well plate screening hits. These findings, thus, pave the way towards the systematic implementation of commercially available FACS for the directed evolution of metalloenzymes making ultrahigh-throughput screening more broadly accessible. The use of microfluidics for the formation of uniform compartments with precise control over reagents and cell encapsulation further facilitates the establishment of highly reliable quantitative assays.

## Introduction

Progresses in protein engineering, DNA sequencing, and bioinformatics, permit the engineering of enzymes to a specific need. Enzymatic properties have evolved over thousands of generations by natural selection. Directed evolution brings forward new tools to accelerate this process, make it more specific or even open the door to develop new-to-nature reactions. Early directed evolution campaigns have resulted in improve solvent tolerance,^[1]^ catalytic rate,^[2]^ selectivity,^[3–5]^ temperature stability, etc.^[6–8]^ Despite these achievements, directed evolution has not yet reached its full potential due to the critical requirement of the preservation of genotype-phenotype linkage during the screening process. Accordingly, most directed evolution campaigns rely on screening libraries in microtiter plates (MTP). Such screening campaigns are time- and resource-intensive and scale exponentially with the number of amino acid positions screened simultaneously.^[9,10]^ These aspects limit the investigation possibilities, constraining scientists either to restrict themselves to the screening of single positions iteratively or to screen only a limited portion of the possible variant landscape.

Within the realm of evolvable enzymes, artificial metalloenzymes (ArMs) consisting of an abiotic metal cofactor anchored within a protein have attracted increased attention recently. Indeed, such hybrid catalysts can endow unique new-to-Nature reactivities within an evolvable protein scaffold.^[11]^ In this context, ArMs based on the biotin-streptavidin technology have been genetically engineered to catalyze a dozen reactions including metathesis,^[12]^ transfer hydrogenation, and hydroamination, hydroxylation, etc.^[13–15]^ Other versatile ArM-scaffolds include: carbonic anhydrase,^[16]^ hemoproteins,^[17,18]^ prolyl oligopeptidase,^[19]^ four-helix bundles^[20]^ or the lactococcal multiresistance regulator.^[21]^ Building on the pioneering work of Meggers and coworkers,^[22–24]^ ArMs that catalyze the allylic deallylation of an allyl-carbamate-protected coumarin were previously screened in 96-well plates.^[25,26]^ The artificial deallylase (ADAse) was optimized using a periplasmic approach by simultaneously screening two positions, to afford a twentyfold improvement in a single round.^[26]^ These results highlight that *i)* ArMs are evolvable for non-natural reactions, *ii)* screening within the periplasm of an *E. coli* is possible and *iii)* screening multiple positions simultaneously increases the chance of identifying beneficial cooperative mutations. However, the lack of high-throughput screening tools restricted these studies to the screening of libraries < 500 mutants.

Advances in microfluidics, and particularly in droplet-based microfluidics, over the past 20 years have led to the development of tools allowing for high-throughput screening of large libraries.^[27,28]^ Such tools are based on the encapsulation of single genetic variants (single cells or DNA molecules) in aqueous compartments (droplets) together with the reaction components. Monodisperse droplets can be produced on-chip at throughputs of several thousand Hertz, resulting in the encapsulation of large libraries within seconds to minutes. Moreover, each droplet, isolated from its surroundings by oil, provides a means of maintaining the phenotype-genotype linkage. Additionally, implementing fluorogenic reactions has enabled further advances for high-throughput sorting, such as on-chip fluorescence-activated droplet sorting (FADS) and have led to the optimization of various enzymes.^[27–33]^ However, the droplet sorting speed is typically lower (∼ a few Hz)^[34]^ than the achievable production rate (∼ several kHz)^[35]^ and decreases even further if the sort requires more than two gates, limiting again the size of accessible libraries. Furthermore, using widely available FACS instrumentation, makes droplet sorting more accessible to a broader scientific community, alleviating on-chip sorting. The latter requires trained staff as it is mostly performed with custom-designed and built equipment and software.

Alternatively, this challenge may be addressed through the use of double emulsion droplets (DEs). In contrast to water-in-oil droplets, i.e. single emulsions, DEs are compatible with commercially-available ultra-fast flow cytometers and fluorescence-activated cell sorters (FACS), available at many academic facilities.^[36–38]^ Previous uses of DEs for directed evolution involved DEs produced in batch and not on a microfluidic device, leading to polydisperse droplets with multiple inner aqueous phase compartments.^[39–42]^ Recent developments, however, allow the on-chip generation of water-in-oil-in-water DE droplets. Analogous to single emulsion droplets, monodisperse DEs can be produced on chip at high throughput and allow compartmentalizing single genetic variants (i.e. genotype) and the corresponding reaction products (i.e. phenotype). DEs can be sorted by FACS in several populations at throughputs comparable to those of the production rate (kHz). Accordingly, this strategy is an attractive tool for high-throughput directed evolution of enzymes.

Here, we describe an ultrahigh-throughput assay based on double-emulsion droplet-microfluidics for the *in vivo* directed evolution of an ArM in DEs. If an average *in vivo* MTP assay requires 6 years to screen around 500’000 variants, assuming one person screens sixteen 96-well plates per week, the presented assay achieves the same results within a week. To do so, DEs produced on-chip are used to compartmentalize single *E. coli* cells, catalyzing the biorthogonal deallylation of the allyl-carbamate protected aminocoumarin **2**, to afford the fluorescent product **3**. The DE population showing the highest product fluorescence intensity (FI) is sorted by FACS and sequenced by NGS to identify variants with improved catalytic activity. First, we validated the method by carrying out a model enrichment. We further validated the accuracy of the method by screening a known 400 double-mutant library.^[26]^

## Results

For the ultrahigh-throughput screening of the ADAse, we envisioned the concept presented in Figure 1a. mNectarine-labeled *E. coli* cells expressing the protein of interest, streptavidin (Sav hereafter), are encapsulated on-chip in DEs. After incubation off-chip, the DEs are subjected to FACS. DEs are sorted for mNectarine fluorescence intensity (mNectarine FI) and coumarin fluorescence intensity (coumarin FI). mNectarine allows the sorting of DEs containing an *E. coli* cell, whereas coumarin serves as a readout for catalytic activity. After the sort, the DEs are ruptured and the plasmid is extracted. The gene of interest is PCR-amplified and analyzed by next-generation sequencing (NGS). The identified hits may be subjected to another round of the assay to iteratively evolve the enzyme of interest.

**Figure 1.**
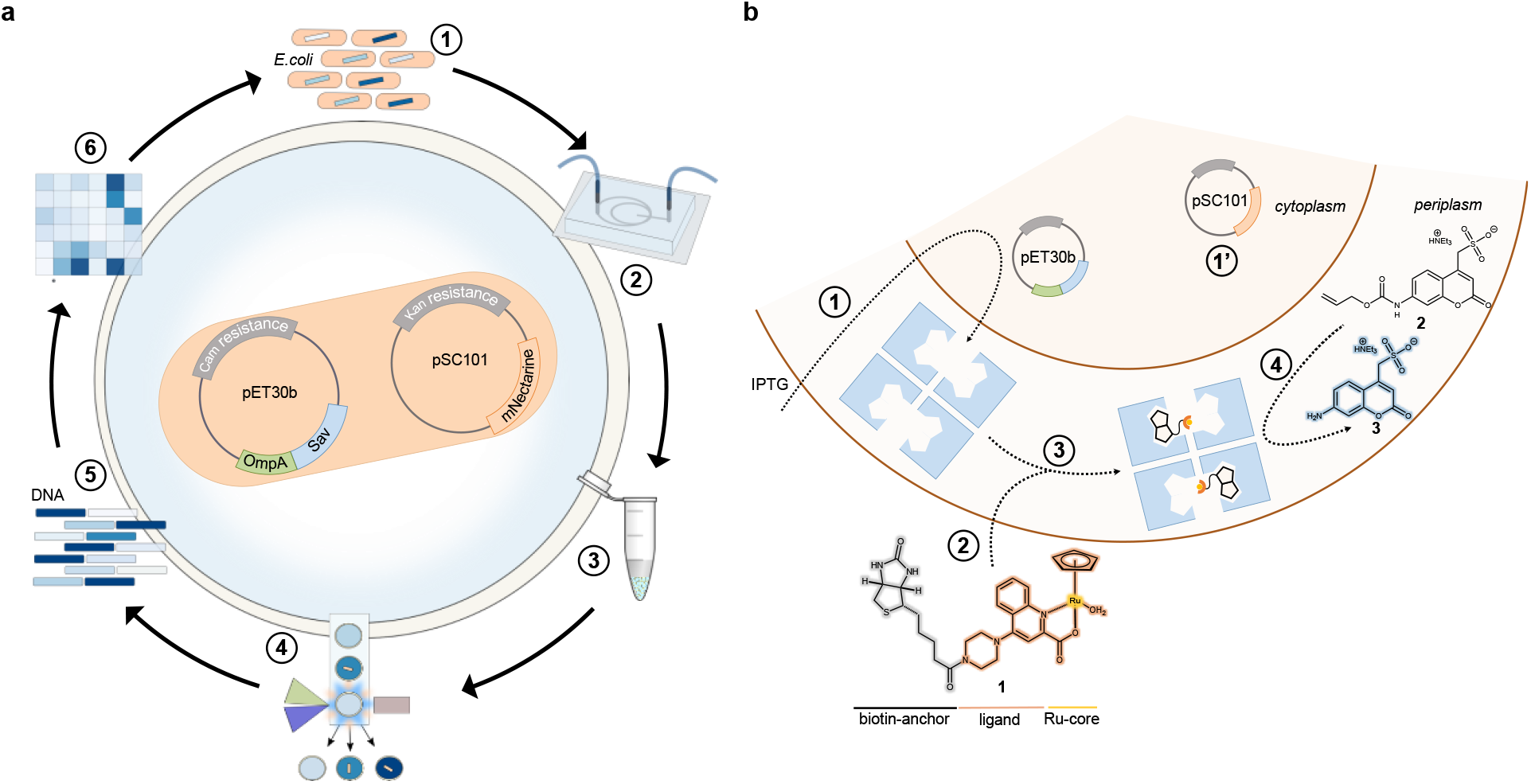
Concept applied for the directed evolution of artificial metalloenzymes using a microfluidics-based screening assay. **a** (1) A library of *E. coli* expressing streptavidin (Sav) in their periplasm and the fluorescent protein mNectarine is generated. (2) The library is encapsulated into double emulsion droplets (DEs) on a microfluidic device together with the substrate **3** and the cofactor **1** and (3) the DEs are incubated off-chip at 37 °C. (4) FACS enables a dual-channel sorting to enrich a population of DEs containing an *E. coli* cell (as highlighted by mNectarine FI, λ_ex_ = 560 nm, λ_em_ = 580 nm) and displaying catalytic activity (as revealed by aminocoumarin **3** FI, λ_ex_ = 405 nm, λ_em_ = 460 nm). (5) Plasmid extraction of the sorted droplets yields plasmid DNA, which is amplified by PCR to enable (6) NGS analysis of the sorted library. Hits can be transformed back into *E. coli* to reiterate the cycle. **b** (1) Sav is encoded on a pET30b vector and induction with IPTG leads to the overexpression and secretion of Sav into the periplasm of *E. coli*. (1’) mNectarine is encoded on a pSC101 vector and is constitutively expressed and compartmentalized in the cytoplasm, enabling the detection by fluorescence of DEs encapsulating an *E. coli* cells. (2) The biotinylated ruthenium cofactor **1** passively diffuses through the outer-membrane into the periplasm. (3) Anchoring cofactor **1** into the binding pocket of Sav affords the artificial deallylase (ADAse). (4) Uncaging of the allyl-carbamate protected substrate **2** yields the fluorescent aminocoumarin **3**, which can be detected and sorted by FACS.

The catalytic system used in this work is the previously described ArM based on Sav compartmentalized in the periplasm and on the biotinylated ruthenium cofactor **1**.^[26]^ The *E. coli* bears two different plasmids: *i)* a pET30b vector encoding a T7-tagged Sav fused to the signal peptide of the outer membrane protein A (OmpA) with kanamycin resistance (Fig. 1b, 1) and *ii)* a pSC101 vector encoding mNectarine with chloramphenicol resistance (Fig. 1b, 1’). The red-fluorescent protein mNectarine is expressed constitutively. Upon addition of IPTG, Sav expression is induced in the cytoplasm, and the protein is subsequently secreted to the periplasm. In the periplasm, Sav forms the homotetrameric protein consisting of four *β*-barrels, which can bind up to four equivalents of biotin.^[12]^ After expression, the cofactor **1** is added and the ADAse is assembled in the periplasm. Pro-fluorescent coumarin **2** is deprotected via an allylic deallylation reaction to afford the fluorescent aminocoumarin **3** which serves as readout for the catalytic activity of the encapsulated variant of the ADAse.

### Characterization of the DE-based screening assay

To implement and optimize the reaction of interest, we developed a microfluidics assay based on the co-encapsulation of single *E. coli* cells together with the substrate **2** and cofactor **1** in water-in-oil-in-water DE droplets. Using a polydimethylsiloxane (PDMS) chip, monodisperse DEs with a diameter of about 15 µm were produced at rates higher than 6000 Hz (Fig. 2a, Supplementary Fig. S1). The *E. coli* sample was incubated with cofactor **1** in PBS and introduced into the chip via a first inlet. The substrate **2**, also dissolved in PBS, was added via a second inlet. The use of two inner aqueous phase inlets provides shielding of the *E. coli* cells and cofactor **1** from the substrate **2** until the on-chip encapsulation. To ensure that only one single *E. coli* cell is encapsulated in a DE droplet, the cell solution was highly diluted, leading to about 20% of DEs containing one cell and 80% of DEs remaining empty. Despite this high dilution, the production rate of 6000 Hz allows the encapsulation of ∼1200 *E. coli* per second, rendering the encapsulation of large libraries feasible: a one million variant library requires less than 15 minutes to encapsulate.^[28]^ After collection and incubation of the DEs off-chip, the fluorescence in the inner aqueous phase was determined using a flow cytometer. DEs were first gated on forward- and side-scatter profile to isolate them from oil droplets and from non-homogeneous DEs, e.g. with two inner aqueous phase droplets instead of one (Fig. 2b). The presence of a fluorescent signal from mNectarine in the DEs reveals the presence of a cell and enables to sort out empty DEs. To assess the coumarin FI and stability, different DE populations were loaded with substrate **2** (500 µM) and three concentrations of product **3** (5 µM, 50 µM and 500 µM). The flow cytometry analysis of these populations 1 h and 24 h after encapsulation confirms that both substrate and product are stably encapsulated in DEs with minimal leakage over 24 h (Fig. 2c). The presence of a sulfonate group on both substrate **2** and product **3** minimizes their diffusion into the hydrophobic oil-phase. Furthermore, these results validate that the three product concentrations can be clearly distinguished from the substrate’s background fluorescence. Similarly, we confirmed that the presence of cells does not affect the substrate integrity nor the catalysis efficiency by co-encapsulating: *i)* cells expressing wildtype streptavidin (wt-Sav) together with 500 µM of substrate **2** (Fig. 2d) and *ii) E. coli* cells without the plasmid for Sav expression together with the substrate **2** and cofactor **1** (Fig. 2e). In both cases, DEs containing cells were compared with DEs without cells just after production and > 10 h later. For condition *i)*, the fluorescence measured in DEs with and without cells is within the same range and remains constant over time. For condition *ii)*, both DE populations display the same fluorescence intensity and behave similarly; their fluorescence increases over time as more substrate **2** is converted into product **3**.

**Figure 2.**
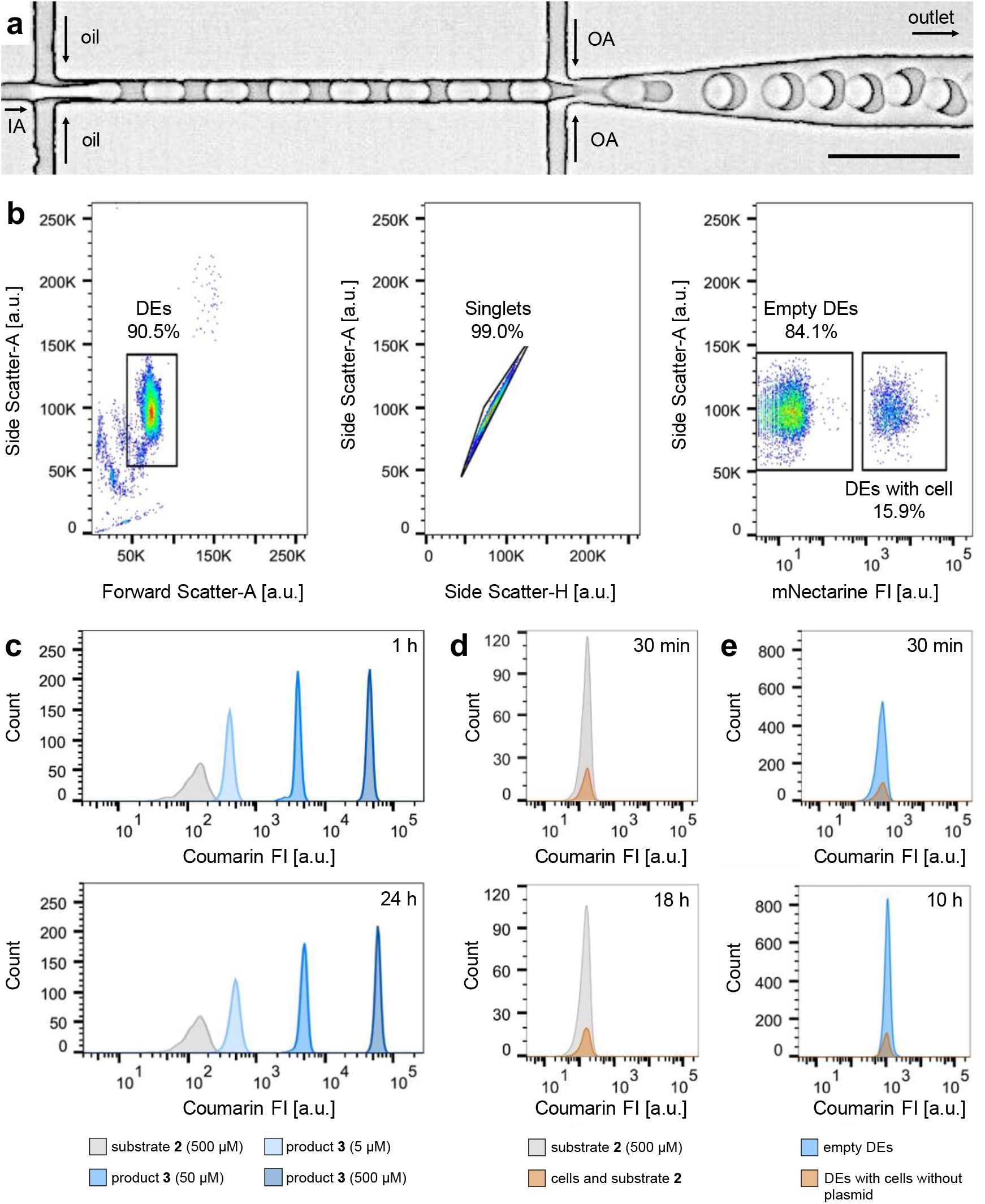
FACS set-up and reaction characterization by fluorescence. **a** Micrograph of the DEs formation. Monodisperse DEs co-encapsulating *E. coli* cells, cofactor **1** and substrate **2** are produced using a PDMS-based microfluidic device (IA: inner aqueous phase, OA: outer aqueous phase, for details see Supplementary Fig. S1). **b** Left: Flow cytometer light scatter gate of a DE sample displaying 10’000 events randomly sampled. Middle: Flow cytometer light scatter gates of homogeneous DEs from the selected DE subpopulation. Right: mNectarine fluorescence intensity (FI) of the selected DE subpopulation allowing the discrimination between DEs containing a cell and empty DEs. **c** Coumarin FI distribution of four different DE samples, encapsulating respectively 500 µM substrate **2** or different product **3** concentrations (5, 50 and 500 µM), obtained 1 h and 24 h after encapsulation. **d** Coumarin FI distribution of DEs encapsulating 500 µM substrate **2** and wt-Sav cells, obtained 30 minutes and 18 hours after encapsulation. **e** Coumarin FI distribution of DEs encapsulating substrate **2**, cofactor **1** and cells without plasmid for Sav expression, obtained 30 minutes and 10 hours after encapsulation.

The screening method was validated by conducting an enrichment experiment in which wt-Sav and a known variant Sav-MR in a ratio 99:1, both expressing mNectarine, were encapsulated in DEs with the substrate and cofactor. The variant Sav-MR bears two mutations at S112M and K121R, and improves the turnover number of the Ru-catalyzed uncaging of the allyl-carbamate protected coumarin **2** to the aminocoumarin **3** (Fig. 3a and 3b).^[26]^ The DE sample was sorted by FACS according to the top 5 % droplets with highest coumarin FI (Fig. 3c). After plasmid extraction, NGS was carried out to determine the enrichment of Sav-MR in the top 5 % gate. Sequence analysis revealed a 68-fold enrichment of Sav-MR over the unsorted sample, thereby validating the reliability of the assay (Fig. 3d).

**Figure 3.**
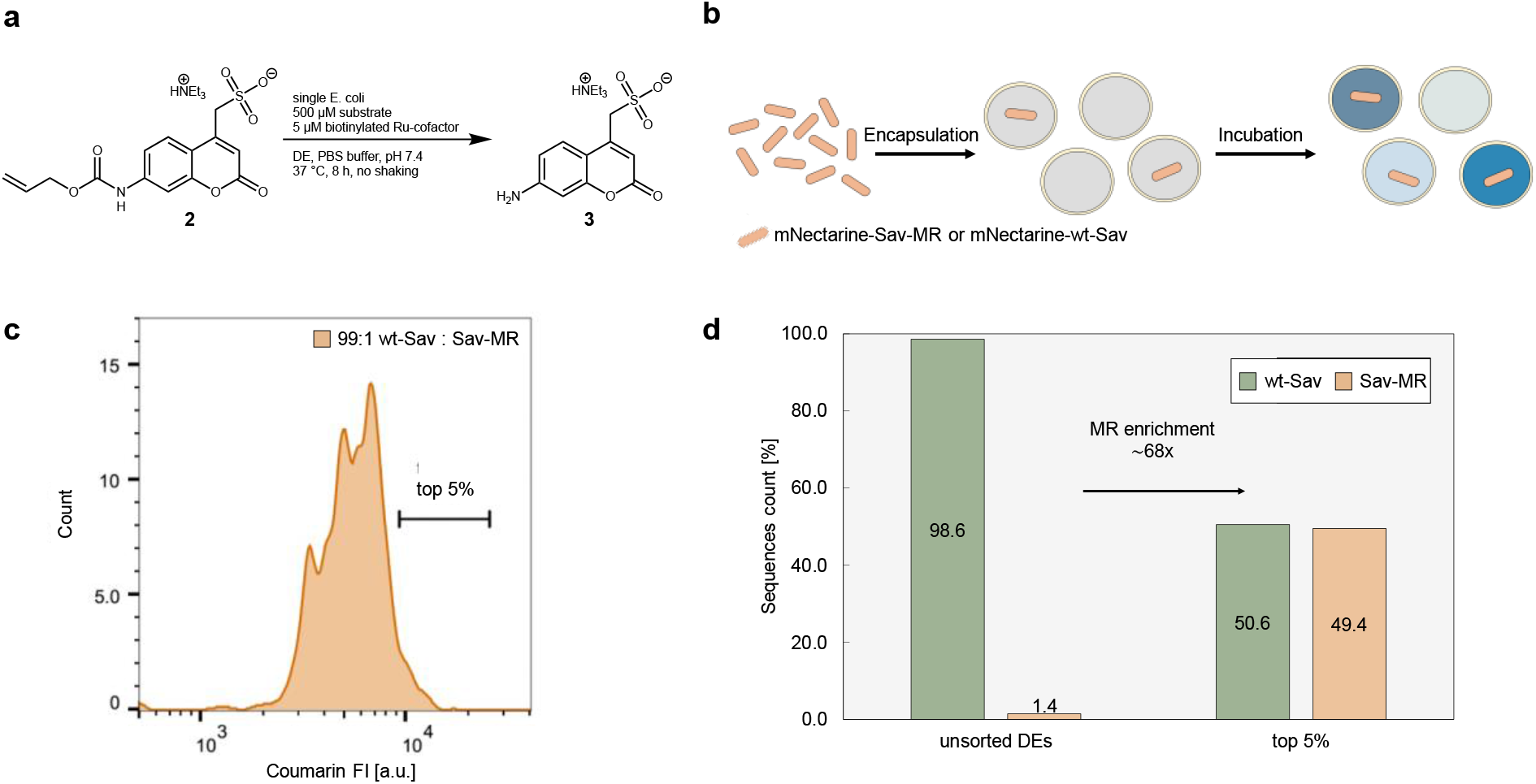
Control experiments with wt-Sav and Sav-MR. **a** Reaction scheme and conditions for the deallylation of the substrate **2** to afford the product **3. b** mNectarine-labeled *E. coli* expressing wt-Sav and Sav-MR at a ratio 99:1 are co-encapsulated with cofactor **1** and substrate **2. c** Coumarin FI distribution of the DEs containing mNectarine-expressing wt-Sav (green) and Sav-MR (orange) in a 99:1 ratio and FACS top 5 % sorting gate. **d** Enrichment of Sav-MR with respect to wt-Sav in the top 5 % of the population determined by NGS.

### Validation of the assay by screening a 400-variant library

The method was further applied to the screening of a known 400-variant library with mutations at positions S112 and K121 of the Sav gene.^[26]^ After preparation of the DE sample and incubation, the top 5 % of the DEs with highest coumarin FI were sorted by FACS (Fig. 4a). Plasmids were extracted from the sorted population and NGS was performed on the top 5 % DE population as well as on the unsorted DEs. The initial study of this library carried out in MTPs by Vornholt *et al*. revealed six highly active variants with activities ≥ 12-fold higher than the activity of wt-Sav: Sav-FQ, -FR, -MR, -MW, -MI and -AW (Fig. 4b, Supplementary Table S1 and S2).^[26]^ Performing the screening in DEs, five out of these six variants were identified by NGS as the variants with the highest increased occurrence in the top 5 % gate compared to their occurrence in the unsorted sample (Fig. 4c and Supplementary Figure S4). Additional variants including Sav-LQ and Sav-MY were found to have higher enrichment in the top 5 % gate over the unsorted sample. These results highlight the validity of the DE screening method and the potential for the time-efficient screening of libraries with minimal reagent consumption. Indeed, with this method the whole screening process, starting with the transformation of the 400-variant library into *E. coli* cells and concluding with the NGS data analysis, was achieved within one week. Moreover, the DE screening led to over 100-fold lower reagent consumption: i.e. 2 nmol cofactor/variant for the 96-well plate screening vs.12.5 pmol cofactor/variant used in the DE screening.

**Figure 4.**
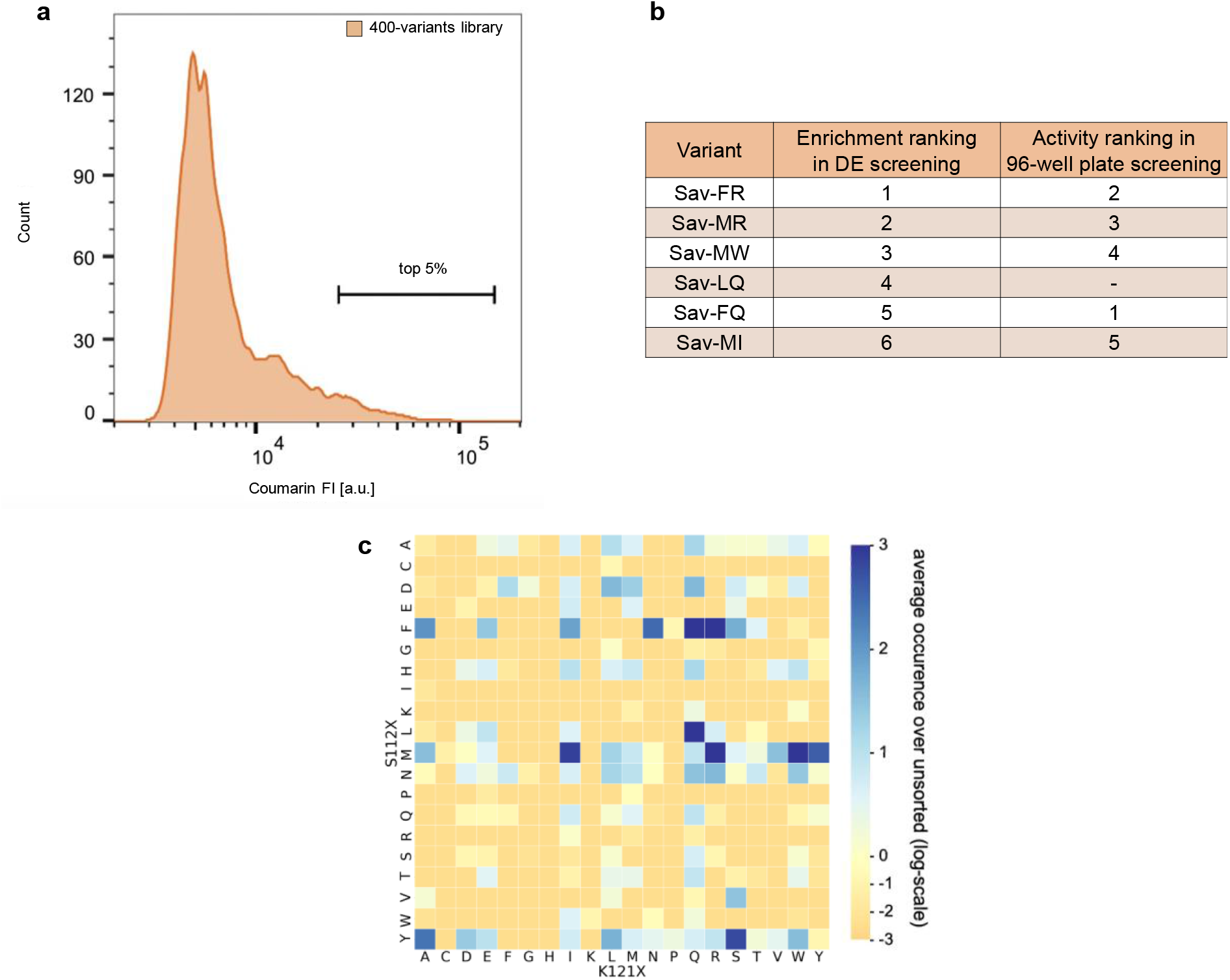
Screening of a 400-variants library in DEs and comparison with a 96-well plate assay. **a** Coumarin FI distribution of DEs containing the library and FACS top 5 % sorting gate. **b** Comparison of the top 6 mutants between DE screening and 96-well plate screening. **c** Enrichment of the 400 mutants over the unsorted sample in the top 5 % of the DE population. x-axis: amino acids at position K121, y-axis: amino acids at position S112, displayed are the normalized enrichment values of the respective mutants over their occurrence in the unsorted sample in a logarithmic scale. Blue: positive enrichment (values over 0), orange: negative enrichment (values below 0).

## Discussion and Conclusion

Capitalizing on droplet microfluidics to encapsulate live *E. coli* at high throughput combined with readily available FACS instrumentation, we developed a straightforward method for rapid screening of ArMs. We exemplified the potential of this method by performing an enrichment experiment of a model library and achieving a > 65-fold enrichment of the desired variant. We further screened a known library of 400 variants within one week using significantly less reagent and successfully identified five out of the six hits found in a previous study.^[26]^ The expansion of this method to the screening of significantly larger libraries (>0.5 Mio variants), which would require over 5 years with standard methods, would not cost significantly more time for the crucial steps that are the encapsulation, incubation and sorting (steps 2-4 in Fig. 1a).

In conclusion our work outlines a powerful method for a streamlined engineering and remodeling of an active site. We foresee that such a screening platform will facilitate and significantly simplify the genetic optimization of (artificial) enzymes.

## Materials and Methods

### Materials

Unless otherwise stated, PBS was purchased from Bioconcept. 1*H*,1*H*,2*H*,2*H*-perfluorooctanol was bought from Fluorochem. Tris-(acetonitrile)-cyclopentadienylruthenium(II)-hexafluorophosphate was purchased from Sigma Aldrich. Water was purified with a Milli-Q-system (Millipore). Antibiotics were purchased from Applichem GmbH. All enzymes, the Monarch plasmid extraction kit, the Monarch PCR and DNA clean-up kit, and the Gibson assembly master mix, were purchased from New England BioLabs. Magnetic beads used for DNA purification were AMPure XP purchased from Beckman-Coulter Life Sciences. Sodium dodecyl sulfate (SDS) was bought from abcr. Poly(dimethylsiloxane) (PDMS, Sylgard 184) was purchased from Dow Corning. Hydrofluoroether (HFE 7500) with 5 % 008-FluoroSurfactant was purchased from RAN Biotechnologies, Inc. All supplies for Nanopore sequencing were purchased from Oxford Nanopore Technologies.

The bacterial strain Top10 DE3 was designed and generously provided by the Panke laboratory.^[12]^ Chemically competent cells (prepared according to the RbCl-method following the Hanahan protocol) and electrocompetent cells bearing the mNectarine plasmid were prepared in-lab.^[43]^

The pET30b vector (Supplementary Fig. S2) was used for the generation of the mutant library. The pSC101 vector (Supplementary Fig. S3) was used for mNectarine.^[26]^ Primers for NGS were designed individually and were synthesized at IDT technologies or Microsynth AG (Supplementary Table S5).

### Methods

For all biological experiments, the equipment was sterilised (121 °C, 20 min). PCR reactions were performed with an Eppendorf Mastercycler Gradient following a general protocol (Supplementary Table S3 and Supplementary Table S4). Agarose and SDS gel electrophoresis chambers and the MicroPulsor Electroporator were purchased from Bio-Rad Laboratories Inc. The gels were visualized with the software Quantity One. DNA concentrations were measured with NanoDrop1000 or the Advanced Analytical 12-capillary Fragment Analyzer from Agilent. Fluorescence scans of individual substances were conducted using a Tecan Infinite M1000 Pro. Large scale protein purification was done using ÄKTA prime Fast Liquid Protein Chromatography System (GE Healthcare). DE droplets were analyzed on a BD LSR Fortessa SORP (Special Order Research Product) and sorted on a BD FACSAria™ II cell sorter. NGS was carried out on an Illumina NextSeq 500 platform.

For the microfluidics assay, neMESYS syringe pumps (Cetoni) and 1 mL syringes (VWR, BD Plastipak Luer-lock) were used to introduce the solutions into the microfluidic chip. We used polytetrafluoroethylen (PTFE) tubing (ID = 0.56 mm, Adtech Polymer Engineering™), precision dispenser needles (23 gauge, Metcal) and metal pins (New England Small Tube, NE-1310-03, 0.025″ OD x .013″ ID x 1.00″ length) to connect the syringes to the chip. To control the chip coating with polyvinyl alcohol (PVA), we used a MFCS-8C pressure control unit (Fluigent).

## Library Preparation

### 400-variants library: pET30b_Sav_library

A glycerol stock containing *E. coli* cells with the 400-mutant library at positions S112 and K121 was grown overnight and the plasmid extracted. The purified plasmid (∼100 ng) was transformed into electrocompetent Top10(DE3) cells (50 µL), containing the plasmid pSC101 with mNectarine encoded. The electroshock was applied using the MicroPulser electroporator by Bio-Rad Laboratories, Inc. Immediately after the electroshock, SOC-medium (450 µL) was added, the reaction transferred to a sterile Eppendorf tube and incubated at 37 °C for 40-60 min. The transformation was split into four equal parts and plated on 12 cm x 12 cm LB-agar plates supplemented with kanamycin and chloramphenicol and incubated overnight at 37 °C. All plates were scraped by the addition of LB-medium (2 mL per plate) and the combined cell suspension was aliquoted. 100 µL cell suspension was mixed with 100 µL of a glycerol stock solution (50%) to obtain 20 aliquots with a ∼25 % final glycerol concentration. The glycerol stocks were immediately frozen in liquid N_2_ and finally stored at -80 °C. One whole such aliquot was used for the inoculation of cultures for the screening assay.

### Substrate and product preparation

Triethylammonium (7-(((allyloxy)carbonyl)amino)-2-oxo-2H-chromen-4-yl)methanesulfonate **2** and triethylammonium (7-amino-2-oxo-2H-chromen-4-yl)methanesulfonate **3** were synthesized as previously reported.^[23,44]^ Stock solutions of substrate **2** (10 mM) and product **3** (10 mM) were prepared in freshly filtered PBS (0.2 µm filter), aliquoted in 60 µL samples to avoid multiple cycles of freezing and thawing and stored at -20 °C for further use.

### Cofactor preparation

The cofactor was prepared *in situ* by mixing a solution of [CpRu(MeCN)_3_]PF_6_ (2 mM in degassed DMF) and a solution of the biotinylated ligand (2 mM in degassed DMF) in a 1:1 ratio inside a glovebox under exclusion of oxygen. The biotinylated ligand was synthesized according to literature.^[25]^ The cofactor **1** stock solution (1 mM in DMF) was used outside the glovebox after incubation of 5-10 min at room temperature.

### General protocol for Sav expression for the screening

A preculture of LB medium (5 mL) supplemented with chloramphenicol (32 mg/mL) and kanamycin (50 mg/mL) was inoculated with the previously prepared glycerol stock of the library of interest and incubated for 8 h at 37°C and 300 rpm. A culture of LB medium (25 mL) supplemented with chloramphenicol (32 mg/mL) and kanamycin (50 mg/mL) in a shaking flask (250 mL) was inoculated with the preculture to a starting OD_600_ =0.05 and incubated for ∼1-2 h at 37°C and 300 rpm (until an OD_600_=0.5-0.8 was reached). Sav expression was induced by the addition of IPTG (50 µM final concentration) and expression was performed overnight at 25 °C and 300 rpm. One sample (1 mL) of cell culture with an OD_600_ = 0.20 was prepared and centrifuged (5 min; 17000 g). The supernatant was discarded and the pellet was resuspended in PBS (990 µL, pH 7.4). To this cell suspension, the cofactor **1** stock solution (10 µL, 1 mM, 10 µM final concentration) was added to afford the cell-cofactor mixture for the droplet production.

## Microfluidic set-up

### Microfluidic platform fabrication

The devices were produced according to a protocol described previously.^[38]^ Briefly, the microfluidic chips were produced using a SU-8 master mold on which a mixture of PDMS and curing agent (ratio 10:1) was poured. The wafer was cured at 80 °C for 3 hours. After punching inlets and outlets with a biopsy puncher (diameter 0.5 mm), the chips were plasma bonded to PDMS-coated glass slides. The device has four inlets for the outer aqueous phase (OA), the oil phase (OP) and the inner aqueous phases (IA1 and IA2), and one outlet. To allow for DE formation, a 2.5 % PVA solution was used to coat hydrophilically the OA and outlet channels according to a protocol previously described by Deshpande *et al*.^[45]^

### Microfluidic assay

DE formation was achieved using flow rates between 0.5 and 5 μL/min for all solutions (IA1, IA2, OA, OP) and was monitored on an inverted microscope (IX71, Olympus) with a high-speed camera (Phantom VEO, Vision Research). In a typical experiment, the solutions contained the following components. OA: PBS with 5 g L^-1^ SDS, OP: HFE 7500 with 2% 008-FluoroSurfactant, IA1: PBS with bacteria (OD_600_=0.2) and cofactor (10 µM) and IA2: PBS with substrate (1000 µM). The DEs were collected in a 1.5 mL Eppendorf tube *via* a short piece of PTFE tubing connected to the outlet.

### DE incubation and FACS

The DEs were incubated statically in collection tubes in an oven at 37 °C for at least 3 hours. For measurements on the flow cytometer, 0.5 µL of DE solution were added to 300 µL of OA (PBS with 5 g/L SDS), agitated manually and loaded on a BD LSR Fortessa SORP (Special Order Research Product). The DEs were gated by size on forward and side scatter profile. The DEs were then gated for singlets using bivariate plots of DEs side scatter height vs area, followed by gating for mNectarine fluorescence (mNectarine: λ_ex_ = 558 nm, λ_em_ = 578 nm; 561 nm laser configuration, bandpass filter 610/20) to determine the presence/absence of cells. Finally, the coumarin fluorescence was determined (coumarin: λ_ex_ = 395 nm, λ_em_ = 460 nm; 405 nm laser configuration, bandpass filter 450/50). For sorting experiments, 10 µL of DE solution were added to 600 µL of OA (PBS with 5 g L^-1^ SDS), agitated manually and loaded on a BD FACSAria™ II cell sorter. This operation was repeated every time the sample tube was empty. During the FACS measurements, the samples were regularly agitated by the operator to prevent the DEs from settling. The DEs were gated as described above and the top 5% of the coumarin **3** peak was sorted into a collection.

## DNA sequencing

### Plasmid recovery

The sorted samples were spun down, and the supernatant (PBS + 0.5 % SDS) was removed. The droplets were then resuspended in the resuspension buffer (B1, 50 µL) of the Monarch plasmid extraction kit. Additionally, perfluorooctanol (12 µL) was added and the suspension was incubated at 50 °C for 20-40 min and 300 rpm. Next, lysis buffer (B2, 50 µL) was added and incubated at room temperature for 1 min. Finally, the neutralization buffer (B3, 100 µL) was added and incubated at room temperature for 2 min. The subsequent steps were carried out following the manufacturer’s protocol (Monarch PCR clean-up kit by NEB), and elution was performed twice with 6 µL of mQ H_2_O (heated to 70 °C).

### NGS measurements

Plasmid DNA (extracted from the five sorted populations and the parent) was amplified using the NGS primers containing the barcodes, adapters and indices for sequencing (Supplementary Table S5). The PCR products were purified by agarose gel electrophoresis and the concentration and purity were determined by capillary electrophoresis. The different samples were pooled according to the measured concentrations to have an equimolar input for the NGS run. The sequencing was carried out with a NextSeq Mid Output v2 kit (300 cycles, PE 2x 150) spiked with additional 20% PhiX. Primary data analyses were performed with Illumina RTA version 2.4.11 and bcl2fastq v2.20.0.422.

## Data evaluation

### Processing of NGS data

We used in-house bash and R scripts to analyze the NGS data. Fastq files containing the forward and reverse reads were obtained following NGS. The reads were extracted and paired. The reads were filtered using the 24-bp fixed region located between position 112 and position 121 (Supplementary Fig. S2), allowing for a maximum of three mismatches. The target fragments were attributed to each sample using their unique barcode (Supplementary Table S5). The mutations at positions 112 and 121 were identified by retrieving the 3 nucleotides flanking prior and post the fixed sequence.

## Acknowledgments

We are grateful to all members of the FACS core facility at the Biozentrum in Basel for the great support and measurements. The Genomics Facility Basel is acknowledged for their support with NGS experiments. Dr. Basilius Sauter is kindly thanked for his support with data analysis and helpful discussions. Further, the authors thank the Swiss National Science Foundation (Grant 200020_182046), the NCCR Molecular Systems Engineering and the European Research Council (Advanced Grant DrEAM - 694424 to T. R. W. and Consolidator Grant HybCell - 681587 to P. S. D.) for their generous support.

