## Supplementary Information for "Single-Round Remodeling of the Active Site of an Artificial Metalloenzyme using an Ultrahigh-Throughput Double Emulsion Screening Assay"

### Affiliations

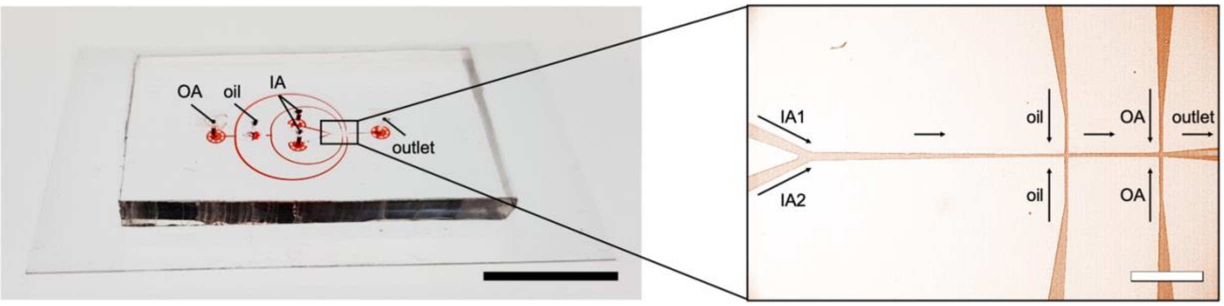


**Supplementary Figure S1. Microfluidic chip for double emulsion production.** Photographs and micrographs of the microfluidic device designed for DE formation. The device has channels with a height of 11 µm and contains 4 inlets: one inlet for the outer aqueous phase (OA), one inlet for the oil (oil) and two inlets for the inner aqueous phases (IA). Black scale bar: 1 cm. White scale bar: 200 µm.

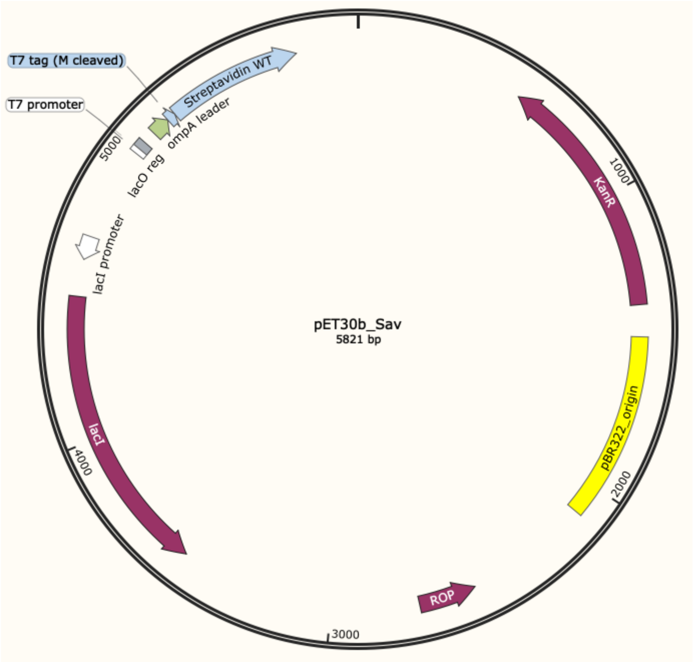


**Supplementary Figure S2. Plasmid map of pET30b encoding Sav.**

**DNA sequence of periplasmic Sav.** red = positions S112 and K121.

ATGAAAAAGACAGCTATCGCGATTGCAGTGGCACTGGCTGGTTTCGCTACCGTA
GCGCAGGCCGCTAGCATGACTGGTGGACAGCAAATGGGTCGGGATCAGGCCGGCAT
CACCGGCACCTGGTACAACCAGCTCGGCTCGACCTTCATCGTGACCGCGGGCGCCGA
CGGCGCCCTGACCGGAACCTACGAGTCGGCCGTCGGCAACGCCGAGAGCCGCTACGT
CCTGACCGGTCGTTACGACAGCGCCCCGGCCACCGACGGCAGCGGCACCGCCCTCGG
TTGGACGGTGGCCTGGAAGAATAACTACCGCAACGCCCACTCCGCGACCACGTGGAG
CGGCCAGTACGTCGGCGGCGCCGAGGCGAGGATCAACACCCAGTGGCTGCTGACCTC
CGGCACCACCGAGGCCAACGCCTGGAAGTCCACGCTGGTCGGCCACGACACCTTCAC

CAAGGTGAAGCCGTCCGCCGCCTCCATCGACGCGGCGAAGAAGGCCGGCGTCAACAACGGCAACCCGCTCGACGCCGTTCAGCAGTAATAA.

**Protein sequence of periplasmic Sav.** red = positions S112 and K121.

MKKTAIAIAVALAGFATVAQAASMTGGQQMGRDQAGITGTWYNQLGSTFIVTAGADGALTGTYESAVGNAESRYVLTGRYDSAPATDGSGTALGWTVAWKNNYRNAHSATTWSGQYVGGAEARINTQWLLTSGTTEANAWKSTLVGHDTFTKVKPSAASIDAAKKAGVNNGNPLDAVQQ.

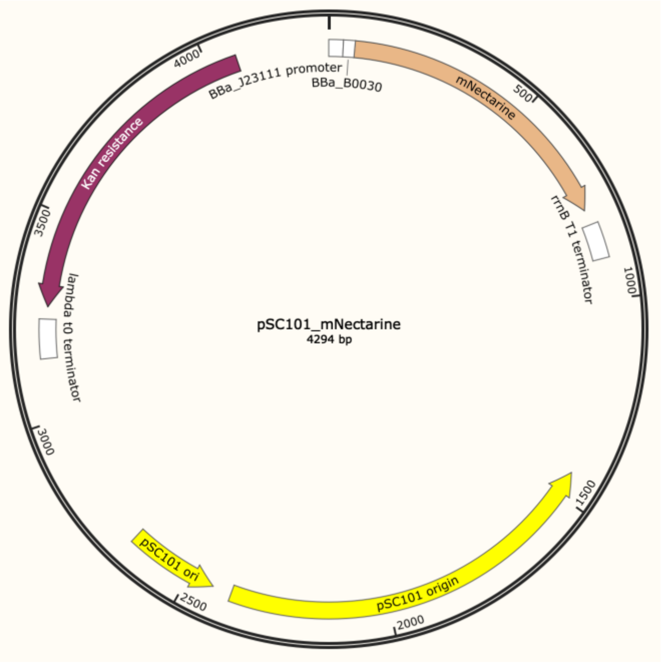


**Supplementary Figure S3. Sequence of constitutively expressed mNectarine.** Plasmid map of pSC101 encoding mNectarine.

**DNA sequence of mNectarine**

ATGGTTAGCAAAGGCGAAGAAGATAACATGGCCATTATCAAGGAGTTTATGCGTTTCAAAGTTCATATGGAGGGTTCCGTCAACGGGCATGAGTTCGAGATTGAAGGCGAGGGGGAGGGCCGTCCGTACGAGGGTACACAAACAGCCAAACTGAAAGTCACGAAGGGTGGACCACTTCCGTTCGCGTGGGATATCCTGTCACCTCAATTCTGCTATGGAAGCAAAGCGTACGTGAAACATCCGGCCGATATTCCGGACTATCTGAAACTGTCGTTCCCTGAAGGTTTAAACTGGGAACGTGTGATGAACTTTGAGGACGGCGGGGTTGTTACCGTAACGCAGGATTCCTCTCTGCAAGACGGCGAGTTCATTTACAAGGTCAAACTCCGTGGTACAAATTTTCCCAGTGATGGTCCAGTTATGCAGTGTCGCACCGTTGGTTGGGAGGCCAGTACCGAACGTATGCATCCGGAGGACGGCGCACTCAAAGGCGAAATCATGCAACGCTT

AAAGCTCAAGGACGGCGGTCATTACGACGCGGAAGTCAAAACAACTTACAAAGCAA
AAAAACCTGTGCAGTTACCGGGTGCATATAACGTCGACATTAAACTCGATATTCTTTC
CCATAACGAGGACTACACAATCGTTGAGCTGTACGAACGCGCGGAAGGCCGTCACA
GCACGGGTGGGATGGACGAGCTCTATAAGTAATAA.

**Protein sequence of mNectarine**

MVSKGEEDNMAIIKEFMRFKVHMEGSVNGHEFEIEGEGEGRPYEGTQTAKLKVTK
GGPLPFAWDILSPQFCYGSKAYVKHPADIPDYLKLSFPEGLNWERVMNFEDGGVVTVTQ
DSSLQDGEFIYKVKLRGTNFPSDGPVMQCRTVGWEASTERMHPEDGALKGEIMQRLKLK
DGGHYDAEVKTTYKAKKPVQLPGAYNVDIKLDILSHNEDYTIVELYERAEGRHSTGGM
DELYK.

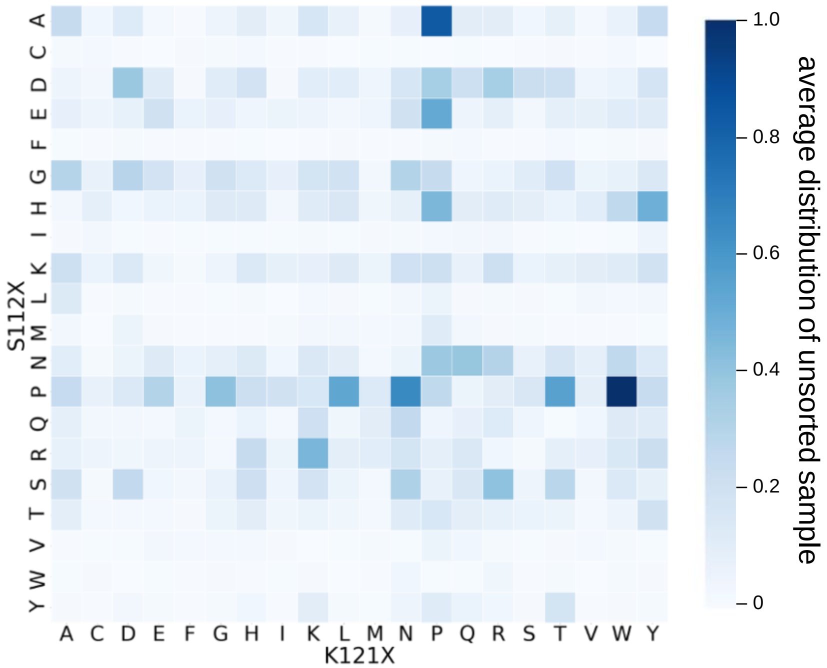


**Supplementary Figure S4. Screening of a 400-variants library in DEs.** Relative distribution of the 400 mutants in the unsorted sample determined by NGS. x-axis: amino acids at position K121, y-axis: amino acids at position S112, displayed are relative occurrences of the respective mutants in relation to the overall reads in the whole sample determined by NGS.

**Supplementary Table S1.** Enrichment factor for the top 20 hits of the DE screening. The enrichment was computed as (count mutantX in top 5 %)/(total count in top 5 %)*(total count in DE)/ (count mutantX in DEs).

| Mutant | DEs count | Top 5% count | Enrichment in top 5% |
|---|---|---|---|
| FR | 416 | 311845 | 741.8 |
| MR | 1072 | 384548 | 355.0 |
| MW | 575 | 71024 | 122.2 |
| LQ | 1201 | 51067 | 42.1 |
| FQ | 1642 | 28326 | 17.1 |
| MI | 1441 | 20822 | 14.3 |
| YS | 1022 | 13524 | 13.1 |
| MY | 834 | 9160 | 10.9 |
| FN | 325 | 3271 | 10.0 |
| YA | 1176 | 10761 | 9.1 |
| FA | 749 | 5136 | 6.8 |
| FI | 153 | 901 | 5.8 |
| FS | 601 | 3106 | 5.1 |
| YL | 1395 | 6892 | 4.9 |
| DL | 8849 | 40409 | 4.5 |
| DQ | 16938 | 76273 | 4.5 |
| NR | 24176 | 108007 | 4.4 |
| YW | 1037 | 4546 | 4.3 |
| MA | 2569 | 10807 | 4.2 |
| MV | 503 | 2093 | 4.1 |

**Supplementary Table S2.** Comparison of the top 20 mutants between 96-well plate experiment (activity measurements) and DE screening (enrichment). Overall, we find a 50% match in the top 10 hits and a 60% match in the top 20 hits.

| **rank** | 96-well plate screening | Activity (improvement over wt-Sav) | DE screening | Enrichment (over unsorted sample) |
|---|---|---|---|---|
| **1** | FQ | 13.8 | FR | 741.8 |
| **2** | FR | 13.4 | MR | 355.0 |
| **3** | MR | 13.4 | MW | 122.2 |
| **4** | MW | 13.3 | LQ | 42.1 |
| **5** | MI | 13.0 | FQ | 17.1 |
| **6** | AW | 12.0 | MI | 14.3 |
| **7** | FS | 11.1 | YS | 13.1 |
| **8** | MM | 10.9 | MY | 10.9 |
| **9** | LR | 10.8 | FN | 10.0 |
| **10** | FT | 10.5 | YA | 9.1 |
| **11** | MV | 10.3 | FA | 6.8 |
| **12** | FA | 10.3 | FI | 5.8 |
| **13** | FN | 10.2 | FS | 5.1 |
| **14** | FE | 10.0 | YL | 4.9 |

| | | | | |
|---|---|---|---|---|
| **15** | YS | 9.7 | DL | 4.5 |
| **16** | IR | 9.5 | DQ | 4.5 |
| **17** | QR | 9.4 | NR | 4.4 |
| **18** | QI | 9.2 | YW | 4.3 |
| **19** | MA | 9.1 | MA | 4.2 |
| **20** | MY | 9.1 | MV | 4.1 |

**Supplementary Table S3.** General reagents set-up for PCR

| Reagents | Appropriate Amounts for one Reaction |
|---|---|
| mQ $H_2O$ | 9.5 µL |
| 2x Q5 maser mix | 12.5 µL |
| Plasmid template (25 ng/µL) | 1.00 µL |
| Forward Primer (10 µM) | 1.00 µL |
| Reverse Primer (10 µM) | 1.00 µL |
| Final volume for one reaction | 25.0 µL |

**Supplementary Table S4.** General PCR program for the library preparation

| Cycle number | Denature | Anneal | Extend |
|---|---|---|---|
| 1 | 95°C, 2 min | | |
| 2-25 | 95°C, 15 s | 65/72 °C, 20 s | 72°C, 2-5 min |
| 27 | | | 72°C, 10 min |
| 28 | | 4 °C | |

**Supplementary Table S5.** Primers used for PCR amplification for Nanopore Sequencing/NGS.

| Primer | Sequence |
|---|---|
| NGS_adapter_L1 | aatgatacggcgaccaccgagatctacactctttccctacacgacgctcttccgatctt**atcacg**agaagcacgcattaataccc |
| NGS_adapter_L2 | aatgatacggcgaccaccgagatctacactctttccctacacgacgctcttccgatctat**cgatgt**agaagcacgcattaataccc |
| NGS_adapter_L3 | aatgatacggcgaccaccgagatctacactctttccctacacgacgctcttccgatctgat**cttgta**agaagcacgcattaataccc |
| NGS_adapter_L4 | aatgatacggcgaccaccgagatctacactctttccctacacgacgctcttccgatctcgat**gccaat**agaagcacgcattaataccc |
| NGS_adapter_L5 | aatgatacggcgaccaccgagatctacactctttccctacacgacgctcttccgatcttcgat**acagtg**agaagcacgcattaataccc |
| NGS_adapter_R1 | caagcagaagacggcatacgagatgtgactggagttcagacgtgtgctcttccgatcttac**gttatt**cggacggcttcaccttg |
| NGS_adapter_R2 | caagcagaagacggcatacgagatgtgactggagttcagacgtgtgctcttccgatctattt**caagaa**cggacggcttcaccttg |
| NGS_adapter_R3 | caagcagaagacggcatacgagatgtgactggagttcagacgtgtgctcttccgatctgatga**tctgcc**cggacggcttcaccttg |
| NGS_adapter_R4 | caagcagaagacggcatacgagatgtgactggagttcagacgtgtgctcttccgatctcgatta**acatag**cggacggcttcaccttg |
| NGS_adapter_R5 | caagcagaagacggcatacgagatgtgactggagttcagacgtgtgctcttccgatcttcgatgc**gtcaac**cggacggcttcaccttg |

blue = i5 indices, violet = i7 indices, orange = barcodes, green = primer binding site.
